# *Pax3* lineage-specific deletion of *Gpr161* is associated with spinal neural tube and craniofacial malformations during embryonic development

**DOI:** 10.1101/2023.07.07.548129

**Authors:** Sung-Eun Kim, Pooja J Chothani, Rehana Shaik, Westley Pollard, Richard H Finnell

**Affiliations:** Department of Pediatrics, Dell Pediatric Research Institute, Dell Medical School, University of Texas at Austin, Austin, TX, 78723, USA; Center for Precision Environmental Health, Department of Molecular and Cellular Biology, Baylor College of Medicine, Houston, TX, 77030, USA; Departments of Molecular and Human Genetics and Medicine, Baylor College of Medicine, Houston, TX, 77030, USA

## Abstract

Shh signaling is the morphogen signaling that regulates embryonic craniofacial and neural tube development. G protein-coupled receptor 161 (Gpr161) is a negative regulator of Shh signaling, and its inactivation in mice results in embryo lethality with craniofacial and neural tube defects (NTDs). However, the structural defects of later embryonic stages in *Gpr161* null mice and cell lineages underlying abnormalities were not well characterized due to their limited lifespan. We found the *Pax3* lineage-specific deletion of *Gpr161* in mice presented with tectal hypertrophy (anterior dorsal neuroepithelium), cranial vault and facial bone hypoplasia (cranial neural crest (CNC)), vertebral abnormalities (somite), and the closed form of spina bifida (posterior dorsal neuroepithelium). In particular, the closed form of spina bifida is partly due to the reduced *Pax3* and *Cdx4* gene expression of the posterior dorsal neural tubes of *Gpr161* mutant embryos involving decreased Wnt signaling whereas Shh signaling was increased. This study provides the novel role of Gpr161 in the posterior neural tube development and confirms its role on CNC- and somite-derived skeletogenesis and midbrain morphogenesis in mice.

## Introduction

The embryonic neural tube is a precursor of the brain and spinal cord, and neural tube development involves coordinated multiple cellular processes and signaling events (Nikolopoulou et al., 2017). The failure of neurulation results in NTDs, a family of congenital malformations that encompass a wide spectrum of phenotypic malformations including anencephaly, spina bifida, craniorachischisis and encephalocele (Ravi et al., 2021; Wilde et al., 2014). Although NTDs are the second most prevalent birth defects in humans, their etiology is not yet thoroughly understood. NTDs are not always isolated malformations, as they are on occasion associated with increased risks for Chiari II malformations (Williams, 2008), Joubert syndrome (Vogel et al., 2012) and Waardenburg syndrome (WS) (Hart and Miriyala, 2017), suggesting the shared genetic etiology and/or pathogenesis of these syndromes with NTDs. In addition, there are several subtypes of spina bifida (SB), including myelomeningocele, meningocele, closed spinal dysmorphisms, and spina bifida occulta (Copp et al., 2015). Some of which are considered to the closed form of SBs due to the skin covered dysmorphism of the spinal cord and resulting from primarily post-neurulation multi-cellular defects. The molecular and genetic pathogenesis of the closed forms of SB are especially not well understood.

Morphogen signaling, such as retinoic acid (RA), sonic hedgehog (Shh), bone morphogenetic protein (BMP), and canonical Wnt/β-catenin signaling, plays a critical role in tissue patterning and organ morphogenesis during embryonic development (Briscoe and Small, 2015). Shh and canonical Wnt/β-catenin signaling regulate dorsoventral and anterior-posterior patterning in craniofacial development and neural tube morphogenesis (Andrews et al., 2019; Le Dreau and Marti, 2012). Specifically, Shh morphogen is secreted from the neuroectoderm of the ventral brain, facial ectoderm, and branchial endoderm and Shh signaling regulates the survival of cranial neural crest cells during craniofacial development (Ahlgren and Bronner-Fraser, 1999; Nasrallah and Golden, 2001). Additionally, Shh morphogen specifies ventral neural identities via being secreted from the floor plate or the notochord, while Wnt morphogen determines dorsal neural identities as they are secreted from the roof plate during neural tube morphogenesis. How they are affected reciprocally at the molecular level despite of their conflicting roles during neural tube development has not well explored. Therefore, any mutations in genes of Shh and Wnt signaling pathway, including *Smoothened* (Jeong et al., 2004), *Suppressor of Fuzed* (*SUFU*) (De Mori et al., 2017; Lu et al., 2014; Svard et al., 2006), and *Protein kinase A* (*PKA*) (Huang et al., 2002; Zhu et al., 2005), are associated with craniofacial and NTDs in both humans and mice.

Pax3 is a paired box motif containing transcription factor that is primarily expressed in the dorsal neuroepithelium of the neural tube, in pre-migratory neural crest cells, and pre-somitic mesoderm in developing embryos (Goulding et al., 1991). Several *Pax3* mutant mice with NTDs, including mutant alleles of *Splotch*, *Sp* (Epstein et al., 1993), *Sp^2H^* (Epstein et al., 1991), *Sp^d^* (Vogan et al., 1993), have been reported. Among them, *Sp* is also considered as an animal model for WS1 with *Sp^Rwa^*(Ohnishi et al., 2017), which also showed NTDs. In addition, *Pax3* mutant mice presented with the cardiac outflow tract septation defects secondary to aberrant cardiac neural crest cell migration and skeletal muscle defects (Franz, 1993), supporting its role in neural crest cell derivatives. In humans, the genetic modification in *PAX3* gene is associated with the etiology of WS and SB (Hol et al., 1995; Nye et al., 1998), suggesting a pathogenic role of Pax3 genes in both diseases.

G protein coupled receptor 161 (Gpr161) is a bona fide negative regulator of Shh signaling in multiple developmental and cellular contexts (Mukhopadhyay et al., 2013). It is primarily localized in the primary cilia for its suppressive role, which activates PKA via increasing cAMP levels, thereby promoting Gli3 processing for the transcriptional inactivation of Shh target genes in the absence of Shh signal. The *Gpr161* null mice are an embryonic lethal by E10.5, and they present with cranial and posterior NTDs with full penetrance, along with facial and limb bud defects. The *Gpr161* hypomorphic mutant mice expressed the spinal NTDs and cataracts (Li et al., 2015; Matteson et al., 2008). Additionally, *Gpr161* conditional mice along with various *Cre* lines, including *Wnt1-Cre*, *Nestin-Cre*, *GFAP-Cre*, and *Prx1-Cre*, showed *Gpr161* depletion mediated developmental defects at later embryonic stages, including intramembranous/endochondral skeletal defects (Hwang et al., 2018; Kim et al., 2021), facial defects (Hwang et al., 2021; Kim et al., 2021), fore-/midbrain abnormalities (Kim et al., 2021; Shimada et al., 2019), ventriculomegaly (Shimada et al., 2019), and limb formation/patterning (Hwang et al., 2018), which are primarily focused on craniofacial and limb defects. In humans, *GPR161* genetic variants were identified in SB patients (Kim et al., 2019) and those with pituitary stalk interruption syndrome (Karaca et al., 2015), respectively. However, the role of Gpr161 in spinal neural tube development remains controversial despite its genetic implication in human SB patients and in *Gpr161* null and hypomorphic SB mouse models.

In this study, we investigated the role of Gpr161 in cranial neural crest and dorsal neural progenitor lineages during mouse embryonic development utilizing *Gpr161* conditional knock out (cKO) mice along with *Pax3-Cre*. We identified that *Gpr161* cKO (*Gpr161^f/f^;Pax3-Cre*) presented with craniofacial defects, cranial vault/facial and vertebrate skeletal defects, and spinal neural tube malformations. In addition, we observed the *Pax3* gene expression was downregulated partly via inhibition of Wnt/μ-catenin signaling in the dorsal spinal neural tubes of *Gpr161* cKO, which could provide the molecular mechanism on Gpr161 mediated spinal neural tube development in mice.

## Results

### *Gpr161* conditional deletion in cranial neural crest and dorsal neural progenitor lineages resulted in the craniofacial defects and spinal malformations

Previous studies demonstrated that *Gpr161* mutant mice present with the craniofacial malformations during mouse embryonic development (Hwang et al., 2021; Kim et al., 2019; Kim et al., 2021; Mukhopadhyay et al., 2013). In addition, *Gpr161* knock out (KO) embryos also presented with delayed posterior neuropore closure (Kim et al., 2019; Mukhopadhyay et al., 2013), which potentially develop into the spinal neural tube malformations at later fetal stages. As *Gpr161* KO embryos are embryo lethal by E10.5, we utilized *Gpr161* conditional knock out mice along with *Pax3-Cre* lines to define the lineage specific role of Gpr161 during craniofacial and spinal neural tube development during later stages of fetal development. For *Pax3-Cre*, Cre was inserted in Pax3 locus (Engleka et al., 2005), and Cre was expressed specifically in the dorsal neural tube, somite, and facial structure in developing mouse embryos. The *flox* control (*Gpr161^f/f^*; *flox* control hereafter) and *Cre* control (*Gpr161^f/+^;Pax3-Cre/+*; *Cre* control hereafter) did not express any phenotypic malformations, whereas *Gpr161* cKO (*Gpr161^f/f^;Pax3-Cre*; refer *Gpr161* cKO hereafter) fetuses showed both craniofacial defects and spinal neural tube malformations (Fig. 1A). We observed these phenotypic malformations with up to 93 % penetrance in *Gpr161* cKO (Fig. 1B and Table 1). In brief, craniofacial defects included tectal hypertrophy, enlarged forebrain and facial defects, initially observed at E13.5, including micro/anophthalmia and microtia, some of which are phenocopied with *Gpr161* cKO with *Wnt1-Cre* in our previous study (Kim et al., 2021). In addition, we observed the spinal dilation starting at E13.5 and various degrees of spina bifida at E17.5 and E18.5 in *Gpr161* cKO fetuses (Fig. 1A and Table 1). There were two representative phenotypic malformations in spinal regions of *Gpr161 cKO* fetuses at E17.5 or E18.5; one showed skin covered without a protrusive sac at the lumbar regions (4 out of 5), while the other affected fetus showed a skin covered lesion with a protrusive sac at the same region (1 out of 5) (Fig 1A). Of note, we could not observe any live born *Gpr161* cKO pups, suggesting *Gpr161* cKO fetuses are embryonic lethal right before birth or possibly during the birth. Less frequently, we observed the kinked tail in *Cre* control mice (Fig. S1), suggesting the potential genetic interaction between *Gpr161* and *Pax3* during mouse tail elongation.

**Figure 1.**
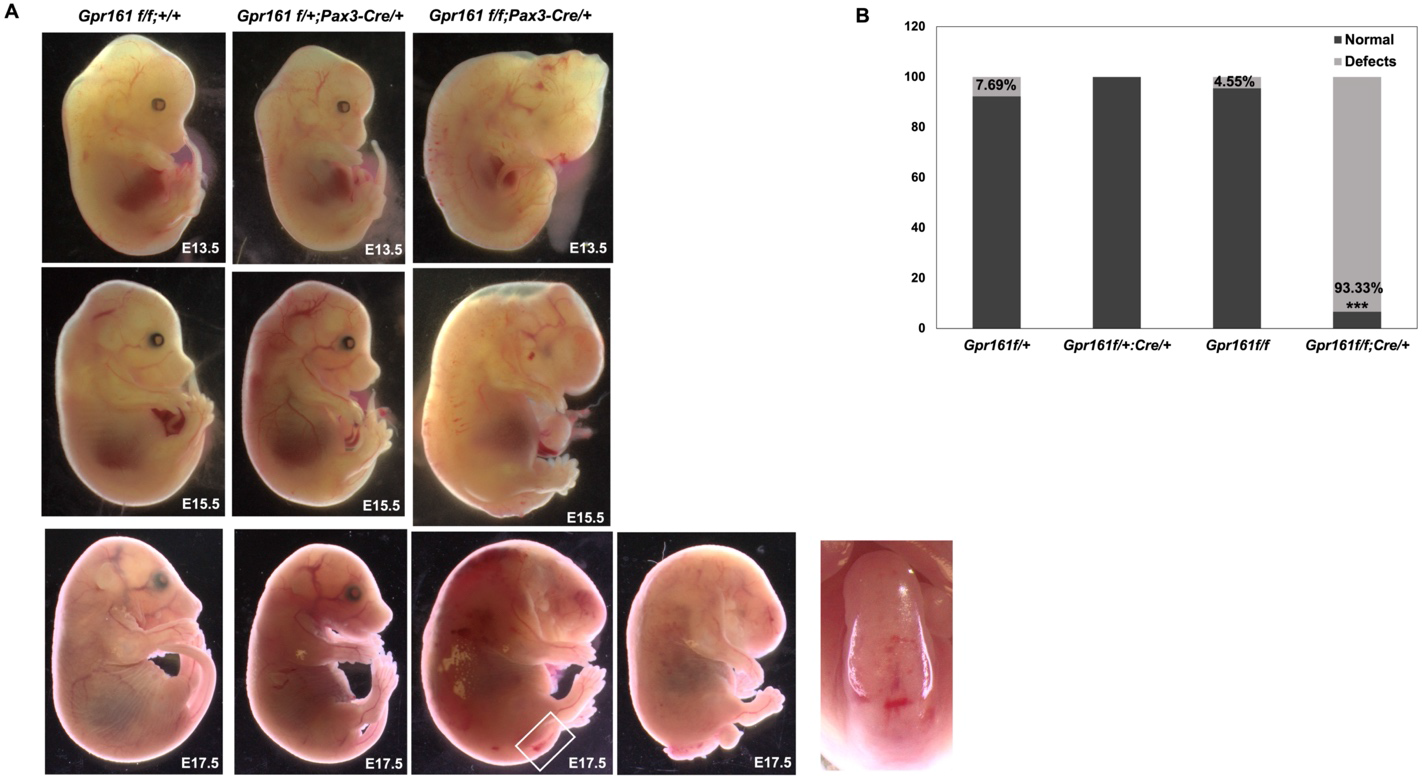
*Gpr161* depletion in *Pax3* lineages results in craniofacial defects and spinal neural tube malformation. (A) Gross morphology of fetuses with *Gpr161^f/f^*, *Gpr161^f/+^;Pax3-Cre/+*, and *Gpr161^f/f^;Pax3-Cre/+* (*Gpr161* cKO) at E13.5, E15.5 and E17.5. The dorsal view of white box in Gpr161 cKO at E17.5 was shown. (B) Statistical analysis based on Table 1. Two sample proportion test was performed between *flox* control vs *Gpr161* cKO and *Cre* control vs *Gpr161* cKO. Both comparisons showed statistically significantly different (***: p<0.001, α<0.05).

**Table 1.**
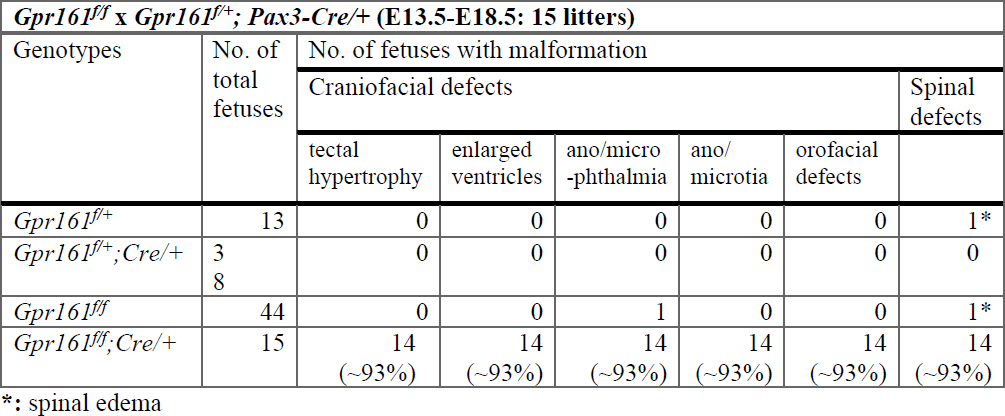
The number of malformed fetuses in *Gpr161^f/f^; Pax3-Cre/+* at E13.5-E18.5.

The histological analysis of anterior and posterior portions of the fetuses were performed to examine the gross structural phenotypic malformations of *Gpr161* cKO (Fig. 2). The tectum and lateral ventricles of *Gpr161* cKO fetuses were extended and the maxilla and mandibles are hypoplastic (Fig. 2A). These facial malformations were phenocopied with *Gpr161* cKO with *Wnt1-Cre* (Kim et al., 2021). At the lumbar level of the spinal cord, the thickness of the neural tubes of *Gpr161* cKO fetuses was significantly reduced compared to WT littermates, and the cystic dilation of the lumens of the spinal neural tube was observed in *Gpr161* cKO fetuses at both E13.5 and E 15.5 (Fig. 2B). The size of the dorsal root ganglia was decreased and abnormally positioned at E13.5 (Fig. 2B). The epidermis layers over the neural tube in the *Gpr161* cKO fetuses were thinner at both E13.5 and E15.5, and the cartilage primordium of vertebrate body was absent at E13.5 (Fig. 2B).

**Figure 2.**
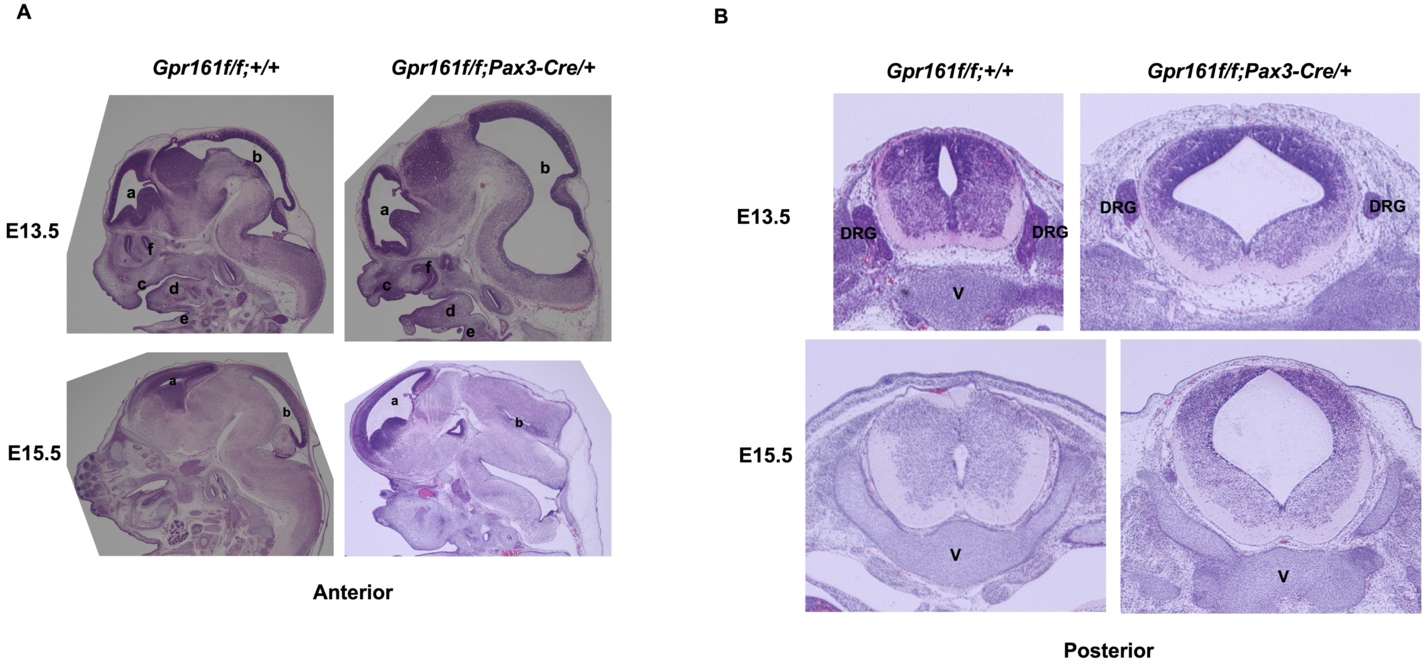
Histological analysis with fetuses of *Gpr161^f/f^* and *Gpr161^f/f^;Pax3-Cre/+* at E13.5 and E15.5. (A) H&E staining of the craniofacies of *flox* control and *Gpr161* cKO fetuses at E13.5 (n=2) (upper) and E15.5 (n=2) (lower). a and b: ventricles, c: maxilla, d: tongue, e: mandible, f: nasal cavity. (B) H&E staining with the posterior portion of *flox* control and *Gpr161* cKO fetuses at E13.5 (n=2) (upper) and E15.5 (n=2) (lower). DRG: dorsal root ganglia, V: vertebrae primordium.

### Gpr161 regulates the craniofacial and vertebrate bone development

Sonic hedgehog signaling is involved in the development of the craniofacial bones via cranial neural crest (CNC) lineages (Jeong et al., 2004), while the *Gpr161* deletion in cranial neural crest lineages resulted in the severe defects in the cranial vault and facial bone formation (Kim et al., 2021). The Cre is expressed in the cranial neural crest lineages as well in *Pax3-Cre* (Engleka et al., 2005) and the gross phenotypic malformations in anterior regions of *Gpr161* cKO fetuses were phenocopied with *Gpr161* cKO with *Wnt1-Cre*. Therefore, we observed the significant reduction of mineralized cartilage and bones derived from cranial neural crest lineages (frontal, maxilla, and mandibles) (Fig. 3A) by performing the Alcian blue (unmineralized cartilage)-Alizarin Red (mineralized cartilage and bones) double staining technique. Much like the *Gpr161* cKO embryos with Wnt1-Cre, the portion of the bone derived mesodermal lineages (parietal bone) was also reduced to lesser degrees. More intriguingly, spinal columns and ribs were significantly misaligned and disorganized along the entire vertebral column from the cervical to sacral regions, and the neural canal was widened in *Gpr161* cKO fetuses at E17.5 (Fig. 3). The vertebral arches were stretched laterally and the cartilage regions between vertebrae are fused in the *Gpr161* cKO fetuses, whereas *flox* control fetuses had well-separated vertebrae (Fig. 3A). To confirm the alterations observed in our skeleton-cartilage double staining procedures, we performed μCT to determine the status of skeletal development and observed the same skeletal malformations including the hypoplasia of craniofacial bones and the disorganized vertebrae formation (Fig. 3B), demonstrating a critical role of Gpr161 in the somite derived vertebral development as well as CNCC derived cranial vault and facial bone development.

**Figure 3.**
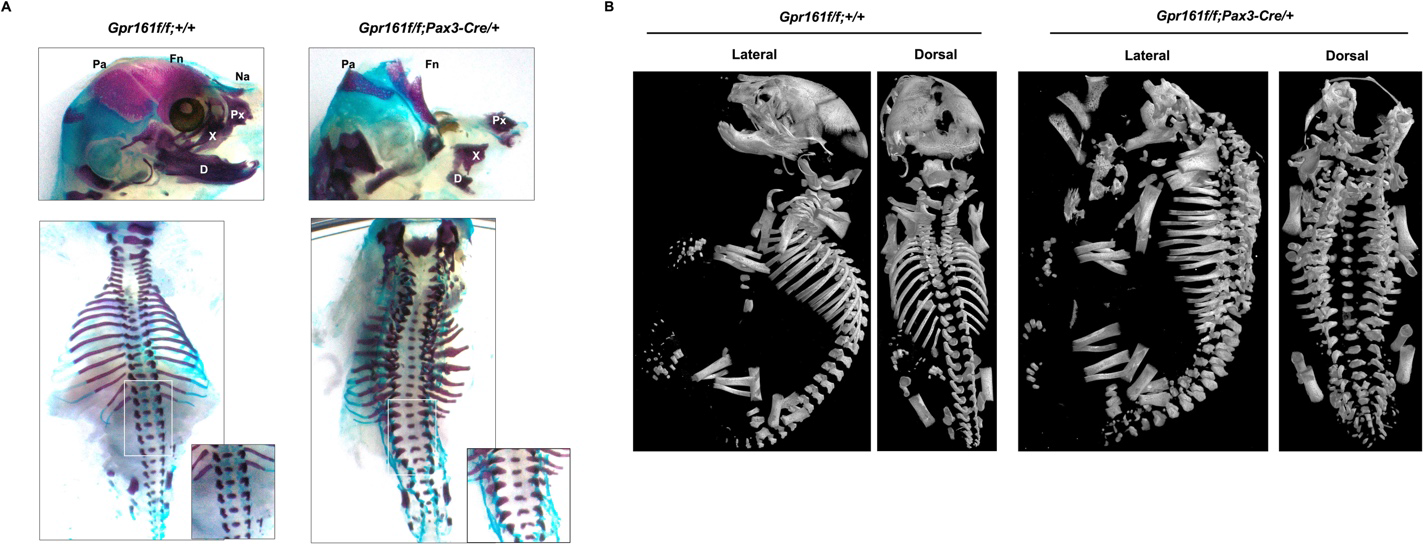
Skeleton analysis in *Gpr161* cKO at E17.5. (A) The skeleton staining of *flox* control (n=3) and *Gpr161* cKO (n=3) at E17.5 with Alcian Blue and Alizarin Red staining. Upper panel: Craniofacial skeletal staining. Fn: frontal bone; Pa: parietal bone; Na: nasal bone; Px: premaxilla; X: maxilla; D: mandible. Lower panel: Axial skeleton staining (B) 3D reconstruction of whole-body skeletons of fetuses from *flox* control (n=3) and *Gpr161* cKO (n=3) at E17.5 with μCT scan. Lateral and dorsal views of fetuses with each genotype.

### The molecular basis of phenotypic malformations in *Gpr161* cKO

In our previous studies (Kim et al., 2019; Kim et al., 2021), we speculated on the molecular basis of craniofacial malformations in *Gpr161* cKO fetuses with *Wnt1-Cre*, which phenocopied the craniofacial phenotypes in *Gpr161* cKO with *Pax3-Cre*. In the present study, we rigorously pursued the molecular basis of the spinal neural tube malformations in *Gpr161* cKO, which is the characteristic phenotypic malformation observed in *Gpr161* cKO with *Pax3-Cre*. First, we determined the *Ptch1* expression in *Gpr161* cKO embryos via whole mount *in situ* hybridization to check the activity of Shh signaling (Mukhopadhyay et al., 2013). *Ptch1* expression was increased overall in *Gpr161* cKO embryos at E10.5 and specifically it was highly expressed in the forebrain, midbrain, frontonasal prominence, first and second branchial arches and in the spinal neural tube (Fig. 4A). Of note, *Ptch1* expression was decreased in the limb buds in *Gpr161* cKO embryos as previously reported (Hwang et al., 2018). Intriguingly, the *Ptch1* expression was significantly increased at the distal extreme of the caudal neural tube in *Gpr161 cKO* embryos (Fig. 4A). *Sox10*, a post-migratory neural crest marker, was expressed in the somites and the branchial arches in both *Cre* control and *Gpr161* cKO embryos without any significant expression changes, whereas *Sox10* expression in the forebrain and frontonasal prominence was slightly decreased (Fig. 4B), indicative of the prominent role of Gpr161 in CNCC migration during craniofacial development. Our previous transcriptomic analysis showed that *Pax3* gene expression was decreased in *Gpr161* KO embryos (Kim et al., 2019), and Pax3 is expressed in the dorsal neural tube during neurulation (Goulding et al., 1991), leading us to examine the involvement of *Pax3* gene regulation. The overall expression of *Pax3* was significantly decreased in *Gpr161* cKO (Fig. 4C) and specifically in the entire dorsal neural tube, somites, forebrain, and the frontonasal prominence, which is negatively correlated with *Ptch1* expression (Figs. 4A and C). We further observed a significant decrease of *Pax3* gene expression in dorsal spinal neural tubes in sectioned *Gpr161* cKO embryos (Fig. 4C; lower panel). In addition, the expression of *Cdx4*, a caudal neural tube marker, was decreased in the tail bud and dorsal neural tube of *Gpr161* cKO embryos (Fig. 4D), providing the role of Gpr161 in the spinal neurulation and tail elongation. To further confirm the Gpr161 mediated regulation of *Pax3* gene expression, we utilized *Gpr161* KO embryos for whole mount *in situ* hybridization. As expected, the *Pax3* expression is significantly decreased in entire *Gpr161* KO embryos compared to heterozygotes (Fig. 4E) specifically, the dorsal side of neural tubes, somites, confirming the regulation of *Pax3* gene expression by Gpr161. Similarly, the expression of *Cdx4* was diminished in the tail bud and dorsal neural tube of spinal neural tube of *Gpr161* KO embryos (Fig. 4F).

**Figure 4.**
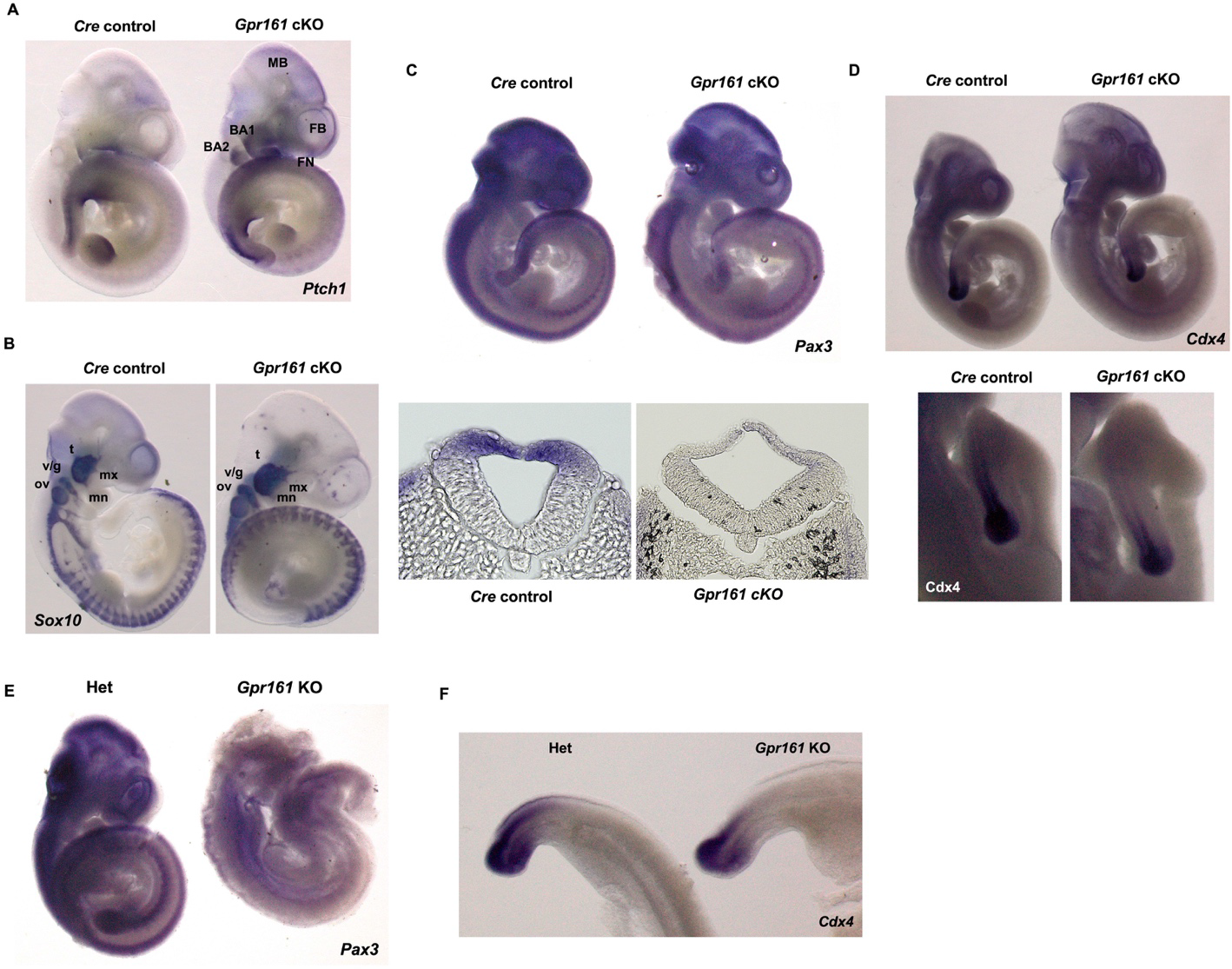
Expression of dorsal/caudal neural tube markers in *Gpr161* mutant embryos. The embryos with *Cre* control and *Gpr161* cKO at E10.5 were used for whole mount *in situ* hybridization with antisense probes against (A) *Ptch1* (n=2), (B) *Sox10* (n=2), (C) *Pax3* (n=3), and (D) *Cdx4* (n=3). The transverse sections from embryos with *Pax3* staining were shown in the lower panel (C). (E and F) The embryos with *Gpr161+/-* (hets) and *Gpr161-/-* (nulls) at E10.5 were used for whole mount *in situ* hybridization with the antisense probes against (E) *Pax3* (n=2) and (F) *Cdx4* (n=2). FN, frontonasal prominence; BA1, branchial arch 1; FB, forbrain; MB, midbrain; trigeminal ganglion; v/g, vestibulo-cochlear/geniculate ganglia; ov, optic vesicle; mn, mandibular branch; mx, maxillary branch.

### Wnt/β-catenin signaling involves in regulating *Pax3* gene expression in *Gpr161* cKO embryos

We were interested in understanding just how *Gpr161* depletion decreased *Pax3* gene expression in the developing mouse spinal neural tube. The previous reports demonstrated that Wnt/β-catenin signaling regulates *Pax3* expression via transcriptional regulation during mouse caudal neural tube closure (Zhao et al., 2014). Our previous study supported the involvement of Gpr161 in the regulation of Wnt/β-catenin signaling during mouse embryonic development (Kim et al., 2019). Therefore, we hypothesized that Gpr161 regulates *Pax3* gene expression via Wnt/β-catenin signaling during caudal neural tube development. To evaluate the Wnt/β-catenin signaling activities in *Gpr161* cKO embryos, we first used Wnt reporter mice (*Tcf/Lef1;H2BB-EGFP*, (Ferrer-Vaquer et al., 2010)) with *Gpr161* cKO. The EGFP signal in the dorsal neural tube of *Gpr161* cKO embryos was significantly decreased (Fig. 5A), suggesting that the Wnt/β-catenin signaling activity was reduced in the dorsal neural tube of *Gpr161* cKO embryos. To further confirm the Gpr161 mediated regulation of canonical Wnt signaling, we observed the expression of *Axin2* gene, a target gene of canonical Wnt signaling, in *Gpr161* KO embryos (Fig. 5B). The overall *Axin2* expression was significantly diminished in *Gpr161* KO embryos, particularly on the dorsal aspect of the neural tube (Fig. 5B; lower panel), demonstrating that canonical Wnt signaling activities were decreased during spinal neural tube development in *Gpr161* KO embryos. In addition, we observed decreased ABC (non-phosphorylated β-catenin), which is another indicator of canonical Wnt signaling activity, in the posterior portion of *Gpr161* cKO embryos (Fig. S2). The overall results support the notion that Wnt signaling activities were decreased in the dorsal side of posterior neural tube in *Gpr161* cKO and *Gpr161* KO embryos.

**Figure 5.**
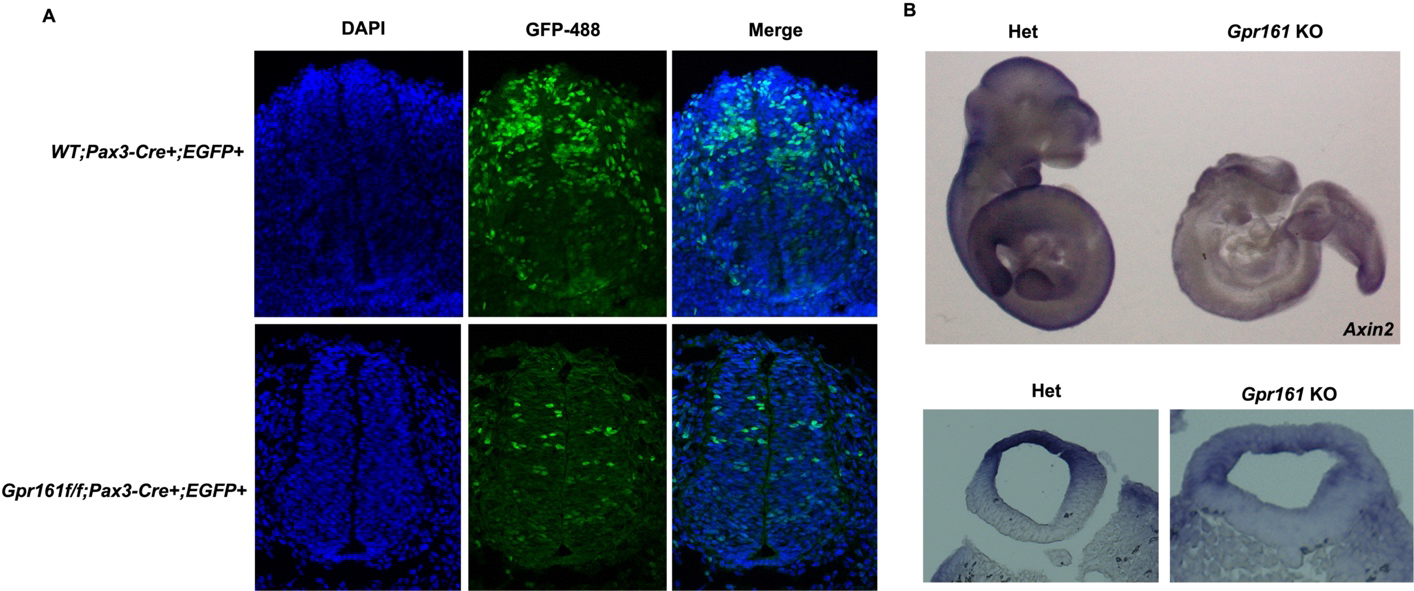
Wnt signaling activities in the caudal neural tubes in *Gpr161* mutant embryos. (A) Immunostaining of cryo-sectioned embryos with *Cre* control (n=2) and *Gpr161* cKO (n=2) harboring the *Tcf/Lef1;H2BB-EGFP* allele. The GFP conjugated with Alexa 488 antibodies was used for EGFP staining and DAPI for staining of the nucleus. (B) The E10.5 embryos of *Gpr161+/-* (het) and *Gpr161-/-* (null) were used for whole mount *in situ* hybridization with *Axin2* probes (n=2). The transverse sections from embryos with *Axin2* staining were shown in the lower panel.

## Discussion

This study revealed the critical role of *Gpr161* in *Pax3* lineages during mouse embryonic development. The *Gpr161* cKO mice presented with two distinct patterns of phenotypic malformations; one involved craniofacial malformation that included tectal hypertrophy and craniofacial skeletal defects, which was phenocopied with *Gpr161* cKO with *Wnt1-Cre* in our previous study (Kim et al., 2021). The second type of altered development involved spinal malformations including spinal neural tube and vertebral skeletal defects, which represents a novel finding in this study. We subsequently observed increased Shh signaling, decreased expression of caudal/dorsal neural tube markers, *Pax3* and *Cdx4* in *Gpr161* cKO embryos, as visualized by whole mount *in situ* hybridization. We further observed the decreased Wnt/β-catenin signaling in dorsal side of the spinal neural tube in *Gpr161* cKO and KO embryos with Wnt reporter mice and by examining *Axin2* expression, respectively, indicating the involvement of Wnt signaling in *Pax3* gene regulation.

### Craniofacial malformations in *Gpr161* cKO

In *Pax3-Cre* mice, Cre is expressed in the dorsal neural tube along the entire anterior-posterior axis and the facial structures at E9 in mouse embryos (Engleka et al., 2005), which is a similar Cre expression pattern as found in the *Wnt1-Cre* mice with respect to the facial structures. Indeed, protrusive tectal defects and the craniofacial skeletal defects were phenocopied in the *Gpr161* cKO with *Wnt1-Cre* embryos. Furthermore, we observed an increased Ki67 staining in the mesencephalic progenitors in *Gpr161* cKO fetuses at E13.5 (Fig. S3A), and the increased *Ptch1* expression in midbrain regions in younger embryos (Fig. 4A), confirming the important role of Shh signaling in midbrain morphogenesis. This was consistent with our previous study with *Gpr161* cKO with *Wnt1-Cre* and *Nestin-Cre* (Kim et al., 2021), and the others with *Ptch1* cKO with *Nestin-Cre* (Martinez et al., 2013). We further identified that the *Pax3* expression was decreased in the dorsal mesencephalon in *Gpr161* cKO as well as *Gpr161* KO embryos at E10.5 (Figs. 4C: upper panels and 4E), providing evidence of the potential regulatory mechanisms of *Pax3* gene expression by Shh signaling during midbrain morphogenesis. This possibility is supported by the previous report (Cairns et al., 2008) that the increased Shh signaling inhibited *Pax3* gene expression with coordination of Wnt/β-catenin signaling during the somitogenesis in chicken embryos. In addition to tectal hypertrophy, the extended lateral ventricles in the forebrain of *Gpr161* cKO at 15.5 was observed (Figs. 1A and 2A), and a similar phenotypic malformation, ventriculomegaly, was described in the *Gpr161* cKO with *Nestin-Cre* (Shimada et al., 2019). What we observed was the increased Shh signaling and the reduced expression of CNCC markers *Sox10* and *Pax3*, in embryonic forebrain at E10.5, providing a strong indication that Shh regulates forebrain morphogenesis via CNCC lineages.

The calvaria and facial bones derived CNCCs were absent or significantly underdeveloped as well as those derived from paraxial mesodermal cells which were also slightly reduced (Fig. 3). These results confirm the role of Gpr161 in the intramembranous ossification during CNCC-derived cranial vault and facial skeletogenesis. It is consistent with ours and others previous observations (Hwang et al., 2018; Kim et al., 2021; Li et al., 2017) that increased Shh signaling in CNCC and mesoderm lineages inhibited calvaria and facial bone formation via dysregulating mesenchymal condensation in the frontonasal and facial prominences. The inhibition of calvaria and facial bone formation was more prominent in *Gpr161* cKO compared to *Sufu* cKO with *Wnt1-Cre*, supporting for the involvement of other signaling pathways during Gpr161 mediated craniofacial skeletogenesis. Indeed, we observed not only increased *Ptch1* expression in the frontonasal prominence and branchial arches of *Gpr161* cKO embryos, but also decreased *Axin1* expression in the same areas of *Gpr161* KO embryos, suggesting the involvement of Wnt/β-catenin signaling in Gpr161 mediated craniofacial skeletal development.

### Spinal malformations in *Gpr161* cKO

The most distinct phenotypic malformation observed in the *Gpr161* cKO embryos compared to *Gpr161* cKO with *Wnt1-Cre* was the spinal neural tube malformation with vertebral defects. The spinal neural tube malformation in *Gpr161* cKO was not a neural tube closure defect, as we could not observe any obvious posterior neuropore (PNP) opening (only less than 5% embryos; Data not shown) right after neurulation (at E10.5), as well as the histological analysis of spinal neural tube of *Gpr161* cKO fetuses (Fig. 2B) did not reveal any sign of an open neural tube, suggesting it could be the closed form of SB. There are discrepant spinal phenotypic malformations between *Gpr161* cKO and *Gpr161* KO; one showed very few open PNPs, while the other has open PNPs with full penetrance at E10.5, respectively. One possible explanation could be the timing and the degree of *Gpr161* depletion, which potentially affects the critically relevant gene expression during spinal neural tube closure. This possibility is supported by the *Pax3* gene expression patterns; while it was significantly decreased in entire embryos, there was a more pronounced reduction in the spinal neural tube in *Gpr161* KO compared to *Gpr161* cKO (Figs. 4C and E) unlike another caudal marker, *Cdx4*, which was equally decreased in embryos of both genotypes. These results further suggest the possibility that Gpr161 is associated with both open and closed SBs via regulating the expression of critical genes in murine spinal neurulation, such as *Pax3*.

The previous report revealed that the transcriptional regulation of *Pax3* by Wnt/β-catenin signaling is critical for β-catenin mediated posterior neural tube closure (Zhao et al., 2014), which lead us to examine the involvement of Wnt/μ-catenin signaling in Gpr161 mediated *Pax3* gene regulation. Our observation in Figure 5, reduced Wnt/μ-catenin signaling activities in the dorsal spinal neural tube in *Gpr161* cKO and KO embryos, suggests Wnt/μ-catenin signaling contributes to a potential regulatory mechanism for Gpr161 mediated *Pax3* gene regulation. In addition, Cairns and coworkers (Cairns et al., 2008) demonstrated that the *Pax3* gene is induced by Wnt that was secreted from the dorsal neural tube and surface ectoderm, and it was repressed with increased Shh gradient via inducing Nkx3.2 during somatogenesis, providing an alternative regulatory mechanism of Gpr161 mediated *Pax3* gene regulation via Shh signaling.

Unlike elevated proliferation in mesencephalon of *Gpr161* cKO, we failed to observe any proliferation changes in the spinal neuroepithelium of *Gpr161* cKO (Fig. S3B), suggesting the involvement of other cellular defects during the spinal neurulation in *Gpr161* cKO. Increased apoptosis was observed in the neural tube of *Splotch* mice (Phelan et al., 1997), which was rescued by inhibition of p53-mediated apoptosis (Pani et al., 2002). In addition, Ptch1 is reported as a pro-apoptotic function in the developing spinal neural tube in mice (Thibert et al., 2003), suggesting the potential role of apoptosis in Gpr161-Pax3-mediated spinal neural tube formation.

The skeletal defects of paraxial mesoderm derived-vertebral columns and ribs in *Gpr161* cKO fetuses (Fig. 3) further suggest the role of Gpr161 in the mesoderm derived endochondral skeletogenesis, which is supported by a previous study with *Prx1-Cre* as the driver (Hwang et al., 2018). The question remains whether the vertebral column defects observed in *Gpr161* cKO fetuses is one of the underlying causes of the closed forms of SB phenotypes.

### Summary and implication to human diseases

This study uncovered the role of Gpr161 in craniofacial morphogenesis and skeletogenesis, as well as spinal neural tube morphogenesis and vertebral formation during mouse embryonic development. We further provide the molecular function of Gpr161 in the regulation of *Pax3* gene expression involving Wnt/μ-catenin signaling during spinal neural tube formation, which partly explains the molecular pathogenesis and genetic association of *Gpr161* in the closed form of spina bifida. Our results suggest a potential molecular target to develop intervention strategies for the prevention of the closed form of SBs with specific genetic mutations. One of which can be the genetic or chemical modulation of Wnt/μ-catenin signaling and/or Pax3 gene to rescue the abnormal phenotypes in *Gpr161* mutant mice, which will be the focus of our future studies.

## Materials and Methods

### Mouse strains and genotyping

All mice were maintained according to the guidelines approved by the Institutional Animal Care and Use Committee (IACUC) of The University of Texas at Austin. *Gpr161* conditional knock out (*Gpr161 flox*) and knock out (*Gpr161* KO) mice were graciously provided by Dr. Saikat Mukhopadhyay (UT Southwestern, Dallas) and the detailed information concerning the generation of transgenic lines was previously reported (Hwang et al., 2018). The transgenic mice, *Pax3-Cre* (#005549) and *Tcf/Lef1;H2BB-EGFP* (#013752), were purchased from Jackson laboratory. The genotypes of the mice and embryos/fetuses were determined by PCR-based genotyping.

### Tissue processing and immunostaining

Embryos were collected at E10.5 from timed mating and processed for cryo-sectioning. To this end, the embryos were fixed, incubated in 30% sucrose solution at 4°C until they were submerged and mounted into optimal cutting temperature (OCT) solution. OCT embedded embryos were cryo-sectioned with 10 μm thickness for immunostaining. The frozen sections were incubated with blocking buffer followed by washing with PBS. Subsequently, they were incubated with GFP-488 (1:200) (A-21311, Thermofisher) in blocking buffer and then with DAPI (1μg/ml) prior to mounting. Images were captured with the aid of a Nikon Ti2E/CSU-W1 spinning disc confocal microscope. For paraffin-embedded sectioning of whole mount *in situ* hybridization embryos, the paraffin-embedded embryos were sectioned at 10 μm thickness, deparaffinized, and the slides were visualized using an Olympus SZX2-ILLT microscope (Olympus, Tokyo, Japan)

### Whole mount *in situ* hybridization

Embryos were collected at E10.5 from timed matings either between *Gpr161 ^f/f^* and *Gpr161^f/+^;Pax3-Cre/+* or between *Gpr161* heterozygotes (*Gpr161+/-*). Collected embryos were fixed and dehydrated with methanol. Whole mount *in situ* hybridization was performed according to standard protocols (Wei et al., 2011). The cDNA plasmids targeting *Pax3* and *Axin2* were obtained from Lee Niswander (University of Colorado), *Sox10* from Yoshihiro Komatsu (UTHealth Houston/McGovern Medical School), and *Ptch1* from Steven Vokes (UT Austin). The DNA template for RNA probes targeting *Cdx4* were generated (Piette et al., 2008) based on the sequence information from the Allen Institute (Primers for DNA templates of RNA probes: *Cdx4*; F-AGTTTACAGGGACCTCAGGATG; R-CAAGAGAAACCAGTGACTCG). Images were captured with an Olympus SZX2-ILLT microscope (Olympus, Tokyo, Japan).

### Bone-cartilage skeletal staining

Skeletal staining was performed using a modified Alcian Blue/Alizarin Red staining procedure (Kessel et al., 1990). Briefly, the E17.5 fetuses were eviscerated and fixed with 95% Ethanol and then Acetone. Fixed fetuses were incubated with staining solution (0,005% Alizarin red S,0,015% Alcian Blue GS in 5% Acetic Acid, 5% H2O and 90% Ethanol) for three days at 37° C. After washing, samples were kept in 1% KOH for 48 hours. For long term storage, specimens serially were transferred into 20%, 50% and 80% glycerol solutions and were ultimately maintained in 100% glycerol. The images were captured with an Olympus SZX2-ILLT microscope (Olympus, Tokyo, Japan).

### Micro-computed tomography and image processing

The E17.5 fetuses were fixed with 10% formalin followed by 70% Ethanol. Specimens were scanned at the University of Texas High-Resolution X-ray CT Facility using the flat panel detector on a Zeiss Xradia 620 Versa. The X-ray source was set to 70kV and 8.5W with no filter. A total of 2001 0.1s projections were acquired over ±180 degrees of rotation with no frame averaging. A source-object distance of 18.0 mm and a detector-object distance of 251.7 mm resulted in 9.98-micron resolution.

## Author contribution

SEK conceived overall study, SEK, PJC, RS, and WP performed mouse experiments and related assays. SEK and RHF wrote and edited the manuscript.

## Acknowledgements

We acknowledge Dr. Saikat Mukhopadhyay (UT Southwestern, Dallas, TX) for providing *Gpr161 flox* and KO mice and Karla Robles-Lopez for the immunohistochemistry work. The H&E staining and the paraffin embedded sectioning of whole mount *in situ* hybridization embryos were done with help of the Histology Core at Dell Pediatric Research Institute and at UT Southwestern, respectively. The μCT images were obtained at the High-Resolution X-ray Computed Tomography Facility of the University of Texas at Austin.

