## Supplementary Material for "*Pax3* lineage-specific deletion of *Gpr161* is associated with spinal neural tube and craniofacial malformations during embryonic development"

**Supplementary Materials**

*Cre* control

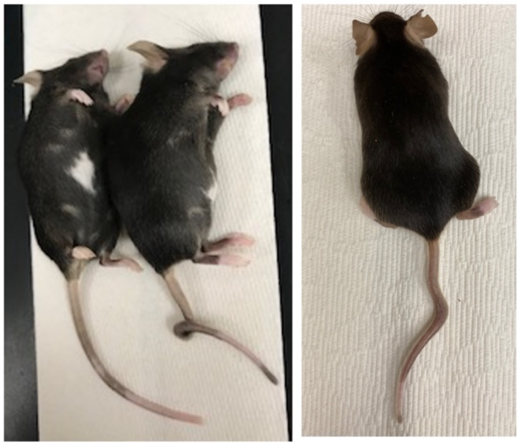

**Supplementary Figure 1. The kinked/curly tail in *Cre* control (*Gpr161*$^{f/+}$*;Pax3-Cre/*+) adult mice.**

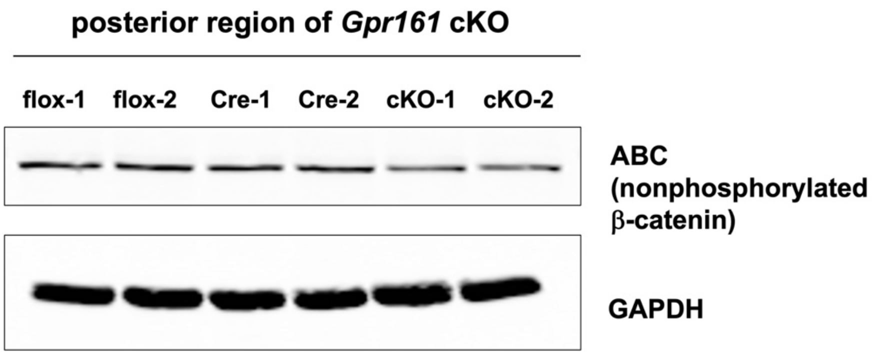


**Supplementary Figure 2. The measurement of Wnt/β-catenin signaling activity in the posterior regions of *Gpr161* cKO embryos at E10.5.** The lysates were prepared from the posterior part of embryos from *flox* control, *Cre* control and *Gpr161* cKO at E10.5, and then performed with Western Blot with antibodies against active β-catenin (ABC) and GAPDH.

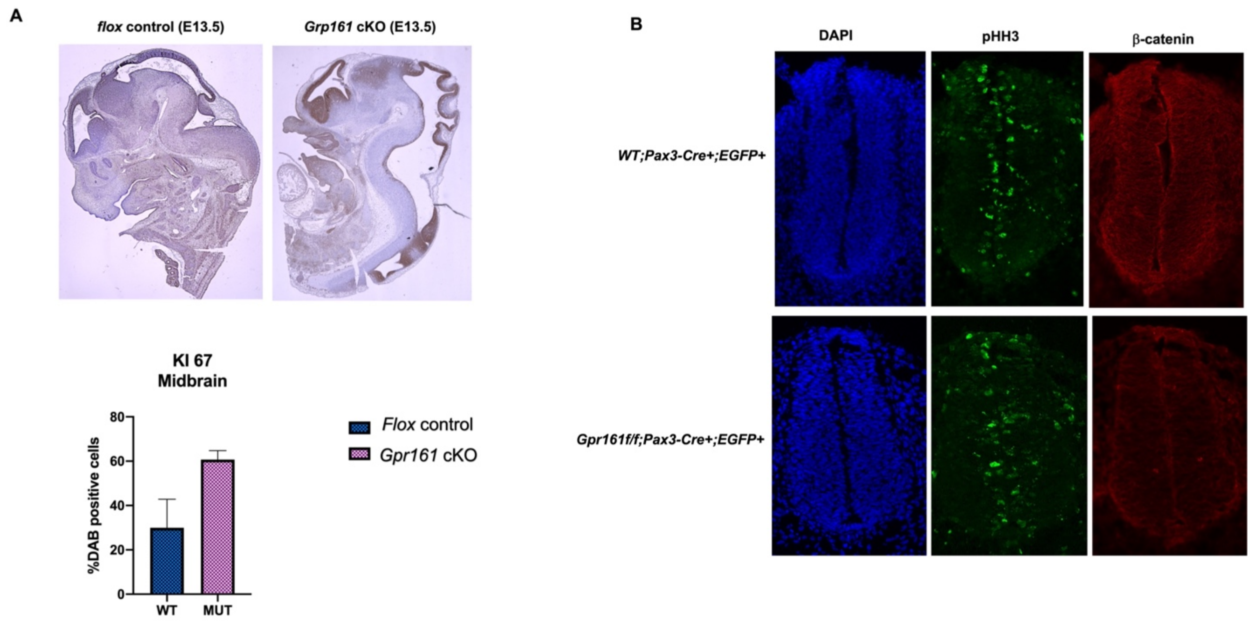


**Supplementary Figure 3. The status of cell proliferation of dorsal midbrain and spinal neural tubes in *Gpr161* cKO.** (A) Paraffin-embedded sections from anteriror part of fetuses of *flox* control (n=2) and *Gpr161* cKO (n=2) at E13.5 were used for IHC. Upper panel: Ki67 staining, Lower panel: Analysis of Ki67 positive cells in dorsal midbrain. (B) Cryo-sections from *WT;Pax3-Cre;Tcf4/Lef1:H2BB-EGFP/+* (n=2) and *$Gpr161^{f/f}$;Pax3-Cre;Tcf4/Lef1:H2BB-EGFP/+* (n=2) were used for immunostaining with phospho Histone H3 (pHH3) and β-catenin. The nucleus was stained with 4′,6-diamidino-2-phenylindole (DAPI).

**Supplementary Material and Method**

**Western blot**

Embryos were harvested at E10.5 from timed mating between *Gpr161* $^{f/f}$ and *Gpr161*$^{f/+}$*;Pax3-Cre/+*. The posterior part of embryos was collected by cutting at the level of hindlimb buds and was lysed with radioimmunoprecipiation assay (RIPA) buffer. The lysates were used for Western blot with active β-catenin and GAPDH antibodies, and then with 1RDye® 800CW goat anti-rabbit IgG (LiCOR). The images were captured by Odyssey® (LI-COR).

**Immunohistochemistry**

Fetuses were harvested at E13.5 from timed mating between *Gpr161* $^{f/f}$ and *Gpr161*$^{f/+}$*;Pax3-Cre/+*. Collected fetuses were fixed, paraffin embedded, and sectioned with 4um-thickness. The paraffin sections were deparaffinized, dehydrated, antigen-retrieved, blocked (blocking solution: Thermofisher scientific), and incubated with primary antibody (Ki67 diluted with Lab Vision$^{TM}$ Antibody Diluent Quanto (Thermo Fischer Scientific) overnight at 4° C. After washing, sections were incubated with Horseradish Peroxidase (HRP) polymer conjugate (UnltraVision$^{TM}$ LP detection system, Thermo fisher scientific) and DAB (Boster bio). The sections were counterstained with hematoxylin (Thermofisher scientific). Images were captured with All-In-One Fluorescent (Keyence) microscope using a 2x and 20x objective. The images were analyzed with Fiji and ImageJ software (NIH).
